# Attracting indigenous pollinators to urban greenspaces

**DOI:** 10.1101/2024.12.17.629021

**Authors:** Luis Mata, Estibaliz Palma, Sally Dawe, Maree Keenan, Phoenix Wolfe, Amy K. Hahs

## Abstract

Urban greening actions across the world are being carried out to support resident, rare and locally extinct insect pollinators. In parallel, an array of best-practise resources have been designed to support build-environment practitioners and professionals seeking to or tasked with implementing greening actions for pollinators. Three main themes permeate these resources: (1) small greenspaces can provide large ecological benefits; (2) high-quality pollinator habitat should include a large share of indigenous plant species; and (3) considering stakeholders’ aspirations, motivations, experience, and concerns is critical for designing effective actions. Despite high interest in implementing these actions, few projects have evaluated their ecological benefits or demonstrated achievement of specific objectives. Even fewer have been specifically co-designed by practitioners and researchers to bridge the science-practice gap. Here, we demonstrate the success of a co-designed greening action in attracting indigenous pollinators to an urban greenspace. We compiled a plant-pollinator interactions dataset in a public park, within a densely urbanised municipality, that was purposefully greened with indigenous plant species to support existing and attract new indigenous pollinators. We then assessed: (1) how pollinator species richness and species-specific occupancy varied amongst plant species, and how these metrics compared between the existing greenspace plant species and the new added indigenous ones; (2) the effect of flower cover of the added indigenous plant species on the probabilities of occurrence of indigenous and introduced pollinators; and (3) the effect of the added indigenous plant species on the structure of the greenspace plant-pollinator ecological network.

An addition of only six indigenous plant species resulted in a 2.5-fold increase in the number of indigenous pollinator species and other flower-visiting insects observed in the park. The added indigenous plants established interactions with all the indigenous pollinators and remarkably almost half of these were found interacting exclusively with the added indigenous plants. Additionally, the number of indigenous pollinators associated with a given plant species was on average 3.5 times higher for the added indigenous plants than for the baseline ones and the probabilities of occurrence of indigenous pollinators were on average invariable higher in the added indigenous than in the baseline plant species. The plant and insect community changes sparked by the greening action resulted in concomitant changes to the structure of the site’s plant-pollinator network. We found an average 4.2-fold increase in the number of interactions linking the greened compared to the baseline network, with almost half of these interactions comprising those between the added indigenous plants and indigenous pollinators. By showing that replacing lawn with indigenous plants leads to positive ecological outcomes for indigenous pollinators and other flower-visiting insects – including increasing the overall greenspace biodiversity by attracting new indigenous insect species to the site – our findings contribute to the evidence base underpinning best-practise resources and provide encouragement to build-environment practitioners and professionals responsible for or endeavouring to support existing or bringing insect pollinators back into our cities. Our study establishes a pathway and acts as a catalyst for researchers and government officials seeking to collaboratively demonstrate that greening is a valuable and sound investment for achieving goals from local, regional and global biodiversity and sustainability plans and policies.

## Introduction

Urbanisation is an acute driver of a wide array of ecological and evolutionary changes experienced by terrestrial and marine organisms living within or near cities (Alberti et al. 2017, Palma et al. 2017, Piano et al. 2017, Johnson and Munshi-South 2017, Merckx et al. 2018, McDonald et al. 2020, Alter et al. 2021, Lambert et al. 2021, Theodorou 2022). Some of the most impactful changes brought about by urbanisation such as habitat modifications, vegetation simplification, microclimatic disruptions, and acoustic, light and chemical pollution may be particularly challenging for insects (Fenoglio et al. 2020, 2021, Tüzün and Stoks 2021, Kotze et al. 2022, Vaz et al. 2023), including pollinators (Harrison and Winfree 2015, Guenat et al. 2019, Baldock 2020, Birdshire et al. 2020, Kurylo et al. 2020, Theodorou et al. 2021, Theodorou 2022, Villalta et al. 2022, Fauviau et al. 2022, Herrmann et al. 2023, Liang et al. 2023, Brasil et al. 2023). Yet, urban environments across the world – from large megalopolis to small villages – may also be a refuge for insect pollinators (Threlfall et al. 2015, Hall et al. 2017, Collado et al. 2019, Nascimento et al. 2020, Zaninotto and Dajoz 2022, Silva et al. 2023) and, when appropriately managed, they can provide opportunities for their conservation (Baldock et al. 2019, Wenzel et al. 2020, Baldock 2020, Braman and Griffin 2022, Brom et al. 2022, Iwasaki and Hogendoorn 2023) and act as hotspots for the pollination services they provide (Theodorou 2022).

Greenspaces within urban areas are attracting widespread interest due to the key role the play in providing food and habitat resources for a diverse array of microbes, fungi, plants and animals (MacGregor-Fors et al. 2016, Threlfall et al. 2017, Spotswood et al. 2021, Fan et al. 2023), with a strong body of literature indicating how greenspace ground and vegetation structure and composition contributes to sustain insect pollinators (Banaszak-Cibicka et al. 2018, Majewska and Altizer 2020, Daniels et al. 2020, Persson et al. 2020, Rajbhandari et al. 2023, Schueller et al. 2023, De Montaigu and Goulson 2024). A core attribute of greenspace vegetation is the quantity and quality of its floral resources, which is of vital importance for the adult stages of bees, butterflies, hoverflies, and other insect pollinators (Hennig and Ghazoul 2012, Baldock et al. 2019, Burdine and McCluney 2019, Wilson and Jamieson 2019, Egerer et al. 2019, Wenzel et al. 2020, Birdshire et al. 2020, Dylewski et al. 2020, Llodrà-Llabrés and Cariñanos 2022). Whilst floral resources such as nectar and pollen may be provided by both introduced and indigenous plant species (Matteson and Langellotto 2011, Baldock et al. 2019, Wenzel et al. 2020, Zaninotto et al. 2023, Howard and Symonds 2023), numerous studies highlight the close ecological associations between locally indigenous plant species and locally indigenous insect pollinator species (Salisbury et al. 2015, Fukase and Simons 2016, Ballare et al. 2019, Buchholz and Kowarik 2019, Cecala and Wilson Rankin 2021, Mata et al. 2021, Anderson et al. 2023, Zaninotto et al. 2023, Abbate et al. 2024, Brown et al. 2024).

A swelling number of urban greening actions are being carried out across the world (Spotswood et al. 2021, Soanes et al. 2023, Gosling 2024), with many exclusively or primarily focused on pollinators (Turo and Gardiner 2019, Braman and Griffin 2022, Tan et al. 2022) and conceptualised to promote the long-term survival of well-established resident pollinators, boost species that have become rare, and bring back locally extinct species (New 2018, Mata et al. 2020). Most projects have been fuelled by a necessity to respond to sustainability and biodiversity policy mandates – for example, the European Union (2020)’s ‘Bringing nature back into our lives’ biodiversity strategy – which has led to an increase in the amount of funding available to national, regional, and local governments to support greening actions for pollinators. Implementing actions for pollinators is also at the heart of wide array of grassroots and non-for-profit community-based organisations (e.g. Pollinator Pathway, USA; The Pollinator Highway, Estonia; B-Lines, UK; Gardens for Wildlife Victoria, Australia; the Melbourne Pollinator Corridor, Australia; Bed and Breakfasts for Birds, Bees, Butterflies and Biodiversity, Australia).

This increase in actions for pollinators has been paralleled by a concomitant surge in the development of a diverse array of best-practise ‘greening for pollinators’ resources, designed to support build-environment practitioners and professionals – from local community champions, school educators, and friends-of-groups volunteers to landscape architects, urban developers, and city planners – seeking to or tasked with implementing greening actions to support pollinators (Coupey et al. 2015, Jordan et al. 2019, Wilk et al. 2021, Mumaw and Mata 2022). Three main themes permeate the body of recommendations provided across these resources: (1) small greenspaces can provide large ecological benefits (Vega and Küffer 2021, Mata et al. 2023); (2) high-quality pollinator habitat should include a large share of indigenous plant species (Salisbury et al. 2015, Ballare et al. 2019, Zaninotto et al. 2023); and (3) considering the aspirations, motivations, experience, and concerns of urban practitioners, stewards, residents, and, most importantly, custodians of Indigenous knowledge, is critical when designing actions (Burr et al. 2016, Mumaw 2017, Turo and Gardiner 2019, Mata et al. 2020, Cumpston et al. 2022, Tan et al. 2022). While interest in planning and executing best-practice actions for pollinators is high, very few projects have sought to design and conduct parallel research to assess the ecological changes brought about by the greening actions and therefore have not been able to provide scientific evidence that specific objectives have been met. Even fewer have been co-designed through partnerships between practitioners and researchers to specifically focus on bridging the science-practice gap (Caron et al. 2014, Cadotte et al. 2017, Kurle 2024).

Here, we demonstrate the success of a co-designed practice-research greening action in attracting indigenous pollinators to an urban greenspace. We compiled a plant-pollinator interactions dataset in a public park, within a densely urbanised municipality, that was purposefully greened with indigenous plant species to support existing and attract new indigenous bee and butterfly species. We then assessed: (1) how pollinator species richness and species-specific occupancy varied amongst the greenspace plant species, and how these ecological metrics compared between the existing greenspace plant species and the new added indigenous ones; (2) the effect of flower cover of the added indigenous plant species on the probability of occurrence of indigenous and introduced pollinator species; and (3) the effect of the added indigenous plant species on the structure of the greenspace plant-pollinator ecological network. Our approach complements traditional analyses focusing on species richness, by harnessing theoretical and analytical advances in ecological community modelling (Kéry and Royle 2016) and network science (Kaiser-Bunbury and Blüthgen 2015, Tylianakis and Morris 2017, Guimarães 2020), as well as by taking advantage of a systems-ecology approach focused on plant-pollinator interactions (Borchardt et al. 2021, Felson and Ellison 2021). As such, our co-designed practice/ research study provides much needed evidence and encouragement that greening actions in urban sites can succeed in driving ecological communities towards robust and resilient states, ultimately protecting and increasing the benefits that nature contributes to people and other species.

## Materials and Methods

### Study site

Our study was conducted at Amersham Reserve, a 17,000 m^2^ greenspace embedded in a dense suburban residential matrix within the City of Greater Dandenong (Melbourne, Victoria, Australia; Fig. S1). The site’s midstorey vegetation prior to August 2020 consisted of a diverse range of groundcover, forb, and shrub species. Of these, 16 species – including species indigenous to the City of Greater Dandenong (5), indigenous to Victoria and/or Australia but not to the City of Greater Dandenong (10) and introduced to Australia (1) – flowered during the study’s data collection period and were included in our plant-insect interactions surveys (henceforth baseline plant species; Table S1). In August 2020, the site experienced a substantial greening action in which a relatively large section of the lawned area of the greenspace was replaced with three garden beds, each spanning approximately 50 m^2^. These beds were planted with a mix of eight indigenous midstorey plant species (henceforth greening action plant species; Table S1; Fig. S2), which were equally spread across each garden bed.

### Plant selection

Plant selection for the garden beds began with a list of species that satisfied the following three criteria set by the municipality’s practitioners: (1) species needed to be indigenous to the City of Greater Dandenong; (2) with moderate to long flowering seasons; and (3) in which indigenous bees (primarily the blue-banded bees *Amegilla asserta* and *Amegilla chlorocyanea*) and butterflies had been previously recorded (Table S2). The full research/practice team then jointly discussed research, management and resourcing considerations, which led to the reduction of the original list to the eight selected species (Table S1). Two of the selected species – white correa *Correa alba* and small crowea *Crowea exalata* – however, did not come to full maturity during the survey period, and were therefore not included in the study.

### Plant-pollinator surveys

We conducted 432 plant-insect interactions surveys across five different timepoints from 25 November 2020 to 4 April 2021. The numbers of days between timepoints varied between 25 and 42 days (mean 33 days). At each timepoint, the interactions between all midstorey plant species (i.e. baseline plus greening action species) that were in flower and pollinators and other flower-visiting insects (henceforth pollinators for brevity) were recorded using a variation of the direct observation survey methodology employed by Mata and colleagues (2020). Each individual plant survey consisted of three time periods of four, three, and three minutes, respectively. During each period, the surveyor actively observed the plant’s flowers, noting down the first sighting of any pollinator that came in touch with the flower’s reproductive organs. Observed pollinators were identified to species/morphospecies on the wing. To complement this direct observation method, the surveyor also photographically recorded as many of the observed interactions as possible using a digital single-lens reflex camera equipped with a 100 mm macro lens. We posteriorly uploaded all photographic records to iNaturalist and benefited from the platform’s community identification systems to check and validate the accuracy of the field identifications. We note that our survey protocol did not include any collection of specimens and therefore no insects were harmed during this study. Surveys were conducted between 10:30 and 17:30 on clear, sunny days with less than 50% cloud cover, and discontinued if rain developed or if wind speed was greater than 5 m/s. For consistency, the species/taxa identification component of all surveys were done by the same researcher (LM). At least four species (tallying less than 5% of all observed interactions) could not be identified beyond order level – these have been recorded as beetles, flies, moths, or wasps, but have not been included in the data analysis.

### Flower cover

To derive the flower cover covariate, we counted the number of flowers (N) within each surveyed plant and measured in mm the length (L) and width (W) of a randomly chosen flower. We then calculated the flower cover (FC) covariate as:

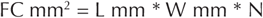

Flower counts varied substantially between the surveyed plants species, ranging from eight – for example, in small lilioids such as the Bulbine lily *Bulbine bulbosa* – to 7,000 – for example, in spreading, groundcover forbs such as the fairy fan-flower *Scaevola aemula*. Similarly, flower size varied markedly between the surveyed plant species, ranging from 0.25 mm^2^ – for example, in some of the surveyed cassinia individuals (genus *Cassinia*) – to 16,900 mm^2^ in some of the surveyed hibiscus individuals (genus *Hibiscus*). While we acknowledge that a more accurate approach would have been to record multiple length and width measurements to capture flower size, the range of plant species included in this study do not show a substantial flower size variation to warrant the extra investment necessary to record multiple measurements.

### Modelling species richness

To assess how pollinator species richness varied amongst the studied plant species, we used a variation of the hierarchical metacommunity model (Kéry and Royle 2016) described by Mata and colleagues (2021, 2023). ‘Plant species’ was the unit of analysis for drawing inferences on pollinator species occupancy and the repeated spatiotemporal samplings constituted the unit of detection replication. The model is organised in three levels: a first level for species occupancy; a second for species detectability; and a third to treat the occupancy and detection of each species as random effects (Kéry and Royle 2016).

The occupancy (Z) and detection (Y) levels were specified as:

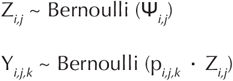

where Ψ_*i,j*_ is the probability that pollinator species *i* occurs at plant species *j* and p_*i,j,k*_ the probability of pollinator species *i* being detected at plant species *j* at spatiotemporal replicate *k*.

The linear predictors were specified on the logit-probability scale as:

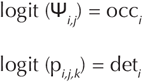

where occ_*i*_ and det_*i*_ are the species-specific random effects, specified as:

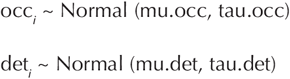

The metacommunity mean occupancy (mu.occ) and detection (mu.det) hyperpriors were specified as Uniform (0, 1) and the metacommunity precision occupancy (tau.occ) and detection (tau.det) hyperpriors as Gamma (0.1, 0.1).

We then used the latent occurrence matrix Z_*ij*_ to estimate the pollinator species richness associated with each plant species SR_*j*_ through the summation:

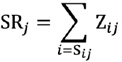

where SR_*j*_ is an indexing vector that accounts for plant-pollinator specificity, that is, for each plant species, the observed pollinator species are included with probability of occurrence = 1 and a limited random subsample of other species occurring in the study area are included with their 0 < Z < 1 estimated probabilities of occurrence (Mata et al. 2021, 2023). This allows us to operate under current understanding of ecological networks’ forbidden links, where non-occurrence of pairwise interactions can be accounted for by spatiotemporal uncoupling, size and reward mismatches, and foraging, physiological and biochemical constraints (Olesen et al. 2010, Jordano 2016).

As these calculations were conducted within a Bayesian inference framework, the pollinator species by plant species estimates were derived with their full associated uncertainties. We then averaged the richness estimates of the (1) baseline plant species and (2) greening action plant species to obtain posterior distributions for each group that could be statistically compared.

### Modelling species-specific occupancy

To assess how the species-specific probabilities of occurrence of indigenous pollinators varied between the baseline and greening action plant species, we used the same model described above but adding a plant group fixed effect (treat_*j*_) to the occupancy level linear predictor:

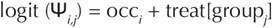

where group_*1*_ and group_*2*_ represent the baseline and greening action plant species, respectively; and treat_*j*_ the plant group fixed effect with priors Normal (0, 0.01).

This analysis was limited to those pollinator species shared between the two plant groups (skippers, grass blue, sweat bees, blue-banded bees and hoverflies).

### Modelling effect of flower cover

To assess the effect of the flower cover on the probability of occurrence of pollinator species, we used a single-species detection-occupancy model (Kéry 2010). ‘Plant species’ was the unit of analysis for drawing inferences on pollinator species occupancy and the repeated sampling periods constituted the unit of detection replication. The model is organised in two levels: a first level for species occupancy and second for species detectability (ref).

The occupancy (Z) and detection (Y) levels were specified as:

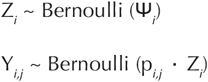

where Ψ_*i*_ is the probability that pollinator species occur at plant species *i* and p_*i,j*_ the probability of the pollinator species being detected at plant species *i* at temporal replicate *j*.

The linear predictors were specified on the logit-probability scale as:

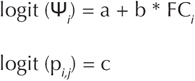

where a and c are the intercept parameters of the occupancy and detection level models, respectively, b the effect (i.e. slope) parameter, and FC_*i*_ the flower cover of plant species *i*. Priors for intercept and slope parameters were specified as Normal (0, 0.1).

We then used the estimated occupancy intercept and effect parameters to draw predictions across a reasonable range of the flower cover gradient (0 – 0.5 m^2^).

This analysis was only conducted for the greening action plant species and independent instances of the model were ran for indigenous and introduced pollinator species.

### Modelling network metrics

To assess the effect of the baseline plant species as compared to the effect of the baseline plus the greening action plant species on the structure of the site’s plant-pollinator ecological network, we first organised the data into plant by pollinator species matrices, with cell values representing the number of times pollinator species were recorded interacting with each plant species. We then used the matrices to calculate: (1) number of interactions; (2) five network-level metrics (linkage density, interaction diversity, interaction evenness, robustness and network specialisation [*H*_*2*_’]); and (3) two species-level metrics (plant [*d*′_plants_] and pollinator specialisation [*d*′_pollinators_]). Metrics were calculated with the R package bipartite (Dormann 2020). Lastly, we used generalised linear models to estimate the group effects on the network metrics. All models were structured around a single level, in which the model for the given metric NM was specified as

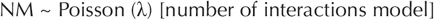

or

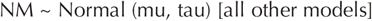

where the expected counts and means were given Normal (0, 0.001) priors and the precision was specified as

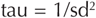

where sd was given a Uniform (0, 100) prior.

A treatment fixed effect treat was added to the linear predictor:

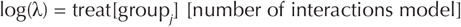

or

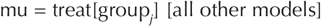

where group_*1*_ represent the baseline plant species and group_*2*_ the combined baseline plus greening action plant species. The treatment group fixed effect treat was given Normal (0, 0.01) priors.

### Bayesian inference implementation

We applied a Bayesian inference approach to estimate model parameters, using Markov Chain Monte Carlo (MCMC) simulations to draw samples from the parameters’ posterior distributions. Models were implemented in JAGS (Plummer 2003), as accessed through the R package jagsUI (Kellner 2024). Three chains of 2,500 iterations were used, discarding the first 500 in each chain as burn-in. We assessed acceptable convergence by visually inspected the MCMC chains and verifying that the values of the Gelman-Rubin statistic fell below 1.1 (Gelman and Hill 2007).

## Results

Overall, we recorded 21 pollinator species including ants, bees and wasps (7), beetles (2), butterflies (6), flies (4) and heteropteran bugs (2). Approximately 70% of the recorded species were locally indigenous to the study area (Table S3), all of which were recorded on the indigenous plant species used in the greening action (Table S4). As much as 47% of these indigenous pollinators were species of bees (1), wasps (1), butterflies (3), flies (1) and heteropteran bugs (1) that were recorded exclusively on the greening action plant species (Table S4). The most commonly occurring species was the introduced European honeybee *Apis mellifera*, accounting for 32% of all records (Table S4). The most commonly occurring indigenous species were woolly sweat bees, accounting for 22% of all records. About half of the pollinator species were recorded six or less times.

### Species richness

The estimated number of indigenous pollinator species found on the greening action plant species was on average 3.5 times higher than that for the baseline plant species (Table S5, Fig. 1). In terms of individual plant species, the sticky everlasting *Xerochrysum viscosum* was the plant species with the highest associated number of indigenous pollinators (Table S5, Fig. 2). The estimated number of indigenous pollinators found on this species was on average 1.3 times higher than that for the next statistically different group – represented by the fairy fan-flower Scaevola aemula and the austral stork’s-bill *Pelargonium australe*, and 2.2 higher than for the other three indigenous plant species (hop goodenia *Goodenia ovata*, cut-leaf daisy *Brachyscome multifida* and bulbine lily *Bulbine bulbosa*) and for the native plant species *Westringia sp. 1*, which was the baseline species showing the highest number of associated indigenous pollinators (Table S5, Fig. 2).

**Figure 1.**
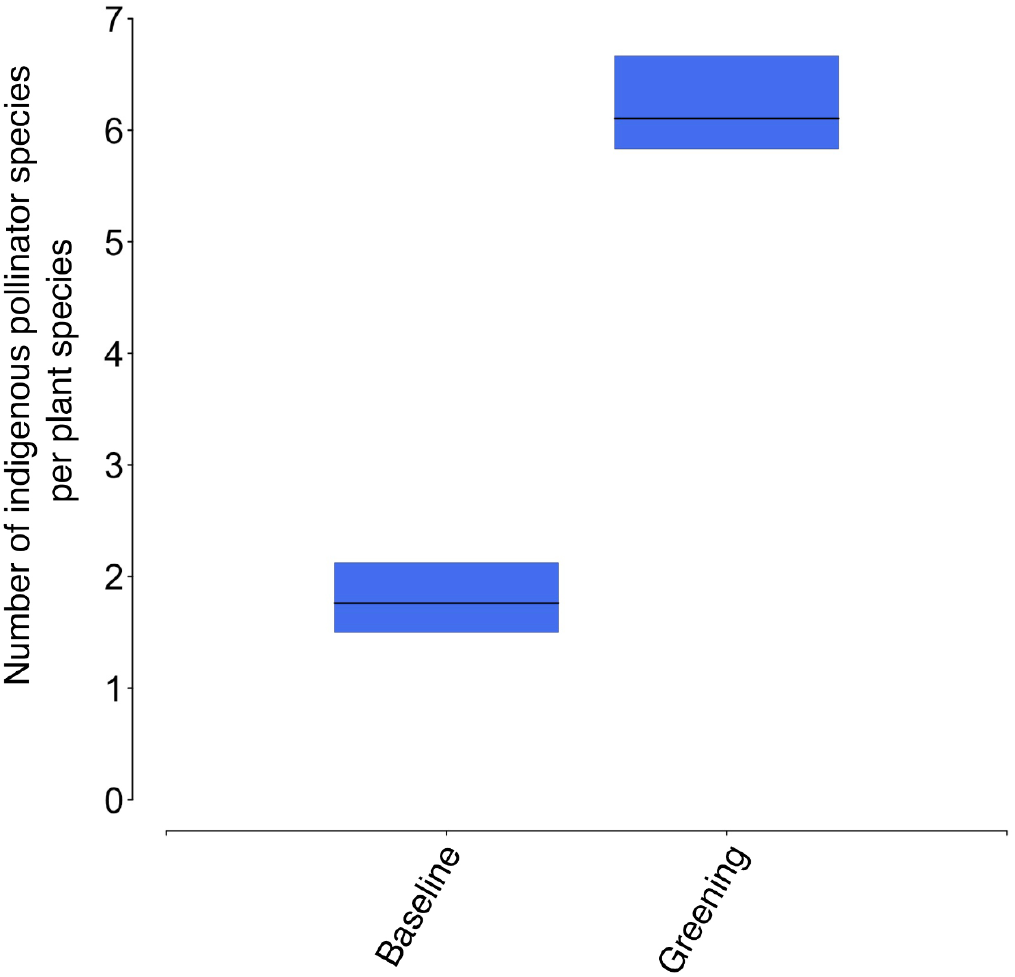
Estimated number of indigenous pollinator and other flower-visiting insect species per plant species for the baseline and greening action plant species. The black horizontal lines represent the mean response and the coloured vertical bars the associated uncertainty (95% credible intervals).

**Figure 2.**
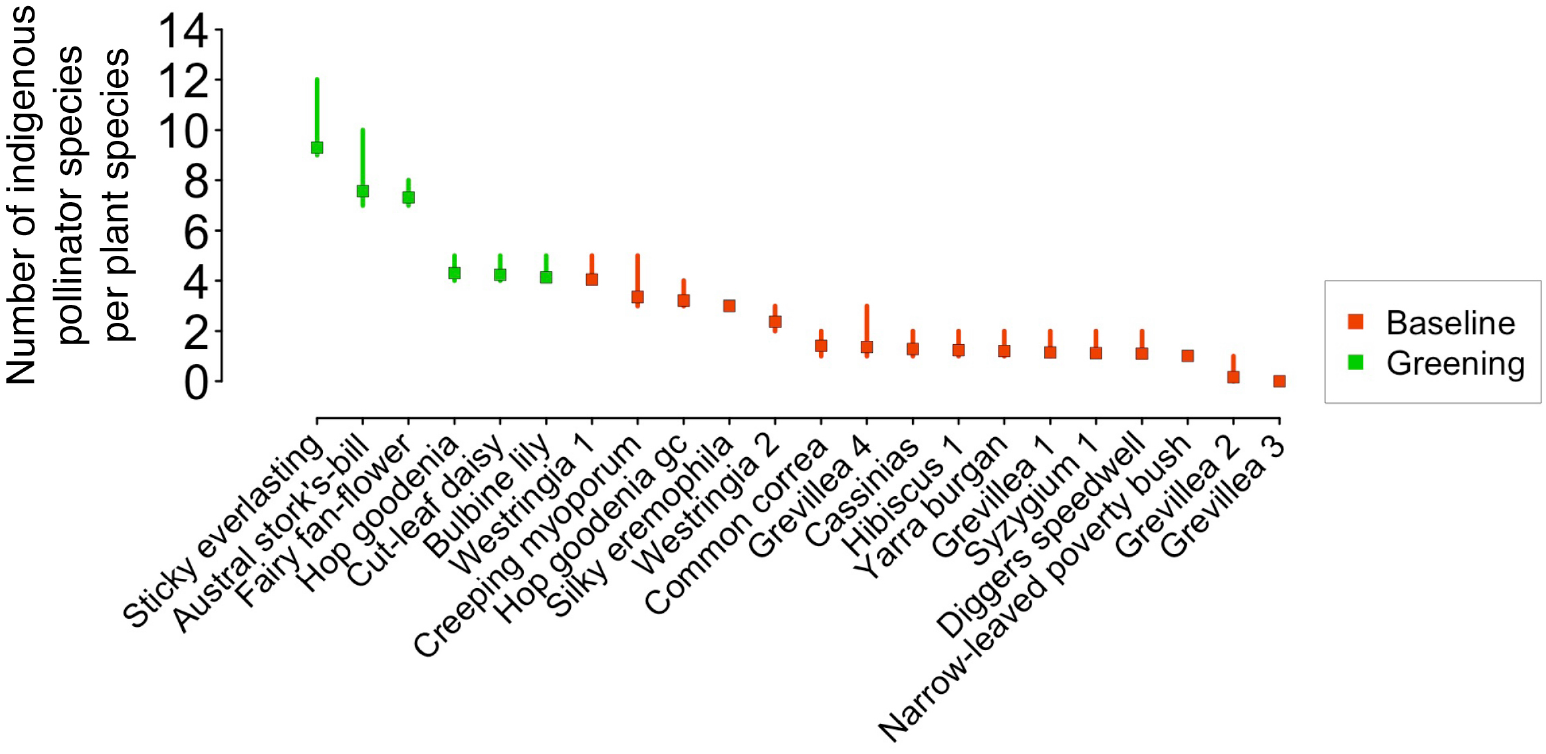
Estimated number of indigenous pollinator and other flower-visiting insect species per plant species for each individual baseline (red) and greening action (green) plant species. The coloured squares represent the mean response and the coloured vertical lines the associated uncertainty (95% credible intervals).

### Species-specific occupancy

The probabilities of occurrence of indigenous pollinator species were on average between 1.7 (woolly sweet bees) and 2.5 (hoverflies) times higher in the greening action than in the baseline plant species (Table S6, Fig. 3). Skippers, blues, and woolly sweet bees showed marginally overlapping 95% credible intervals, while blue-banded bees and hoverflies showed distinctly overlapping 95% credible intervals (Table S6, Fig. 3). The probabilities of occurrence of the two introduced pollinators were on average 1.7 (European honeybee; non-overlapping 95% CI) and 2.2 (cabbage white; overlapping 95% CI) times higher in the greening action than in the baseline plant species (Table S6).

**Figure 3.**
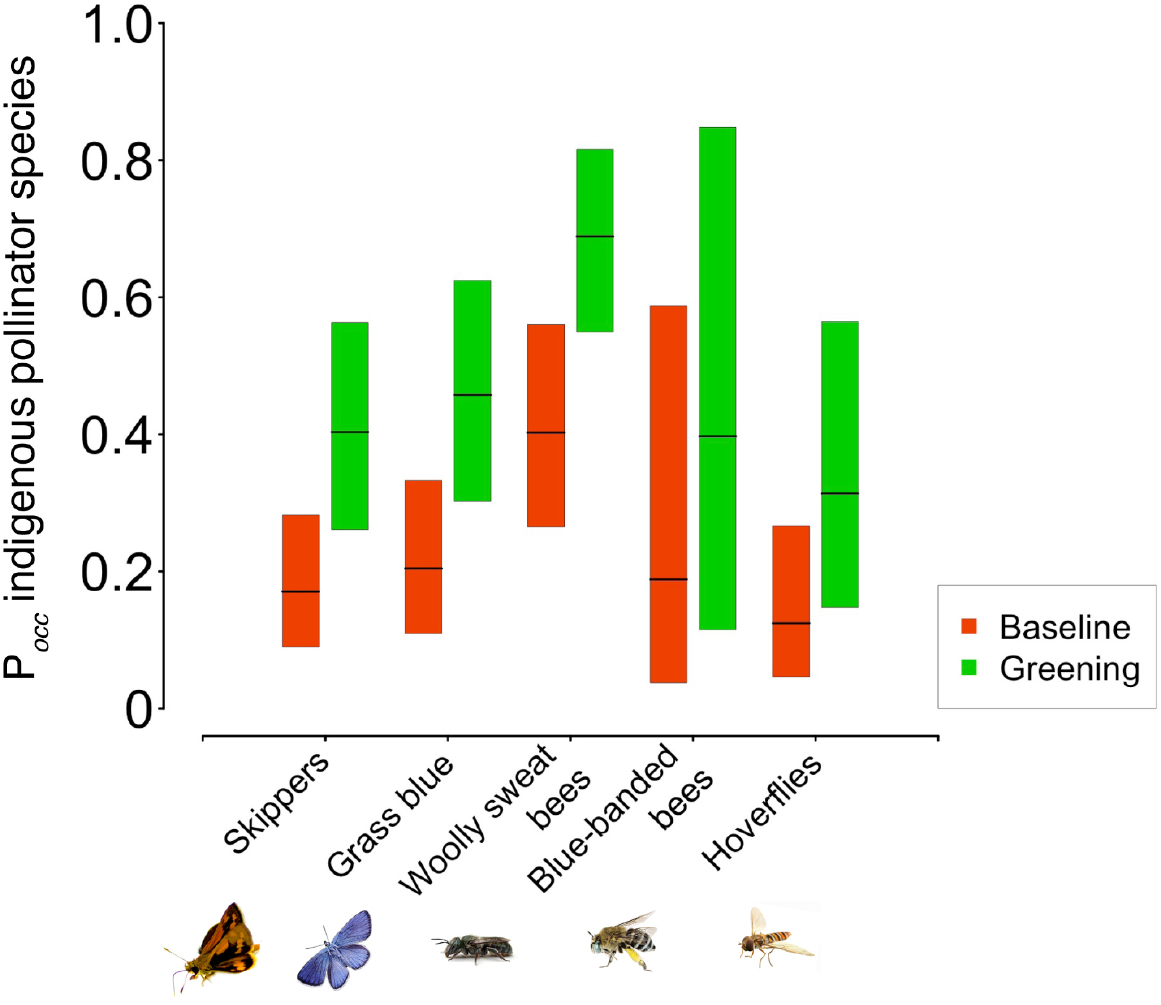
Estimated probabilities of occurrence for indigenous pollinators (skippers, grass blues, woolly sweat bees, blue-banded bees and hoverflies) for the baseline (red) and greening action (green) plant species. The black horizontal lines represent the mean response and the coloured vertical bars the associated uncertainty (95% credible intervals).

### Effect of flower cover

Flower cover had a strong positive effect on the probability of occurrence of indigenous pollinator species (Table S7; Fig. 4). The mean probability of occurrence of indigenous pollinators reached and remained steadily at one in plants species with flower covers higher than 0.2 m2 (Fig. 4). The effect of flower cover on the probability of occurrence of introduced pollinators was also positive but slightly less pronounced (Table S7; Fig. S3).

**Figure 4.**
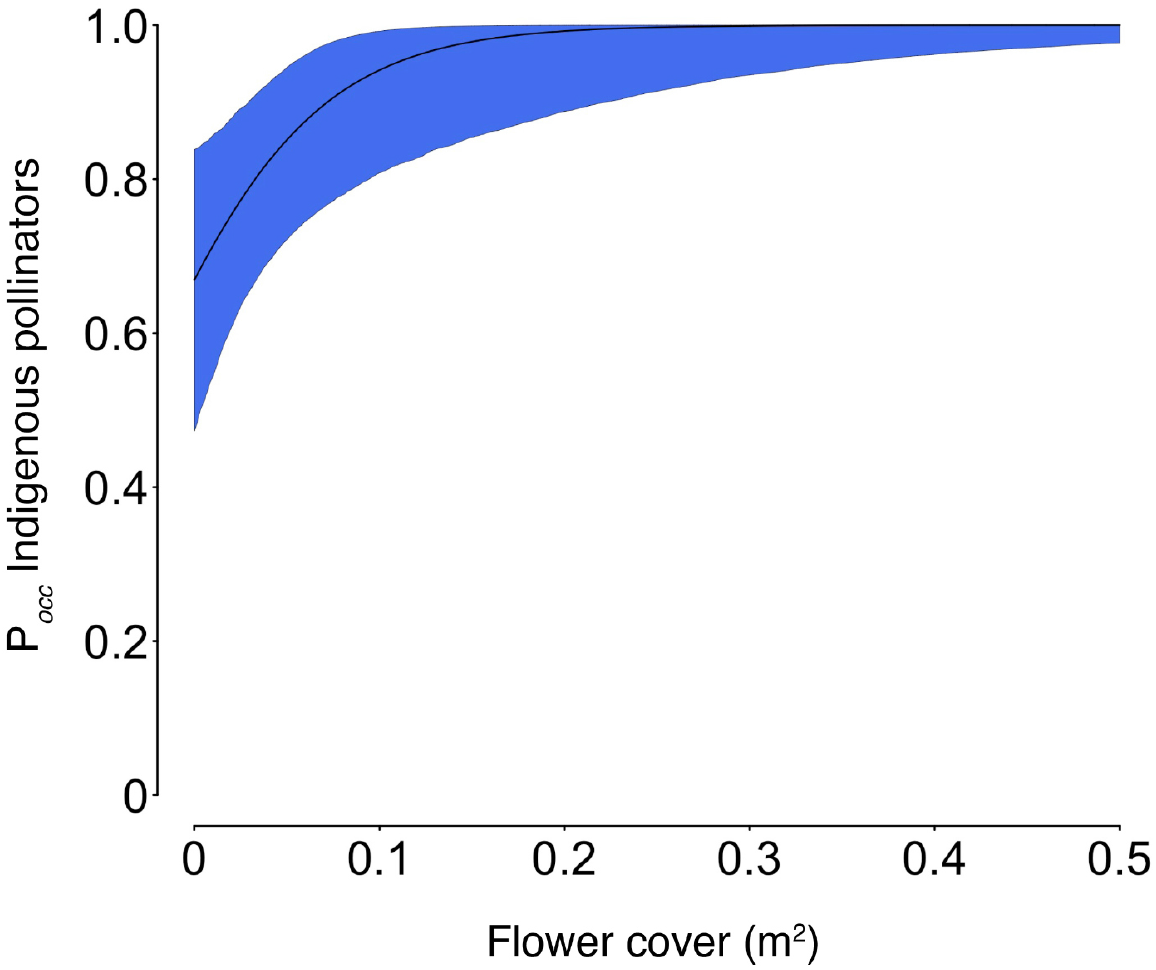
Estimated effect of flower cover on the probability of occurrence for indigenous pollinator and other flower-visiting insect species. The black curved line represents the mean response and the coloured curved polygon the associated uncertainty (95% credible intervals).

### Network metrics

The addition of the greening action plant species caused pronounced changes in the structure of the site’s plant-pollinator network (Fig. 5). The number of interactions was on average 4.2 time higher in the greened compared to the baseline network (Table S8; Fig. 6). Similarly, linkage density and interaction diversity were one average 1.9 and 1.5 times higher, respectively, in the greened compared to the baseline network (Table S8; Fig. 6). Interaction evenness was relatively high across both networks, showing distinctly overlapping 95% credible intervals (Table S8, Fig. 6).

**Figure 5.**
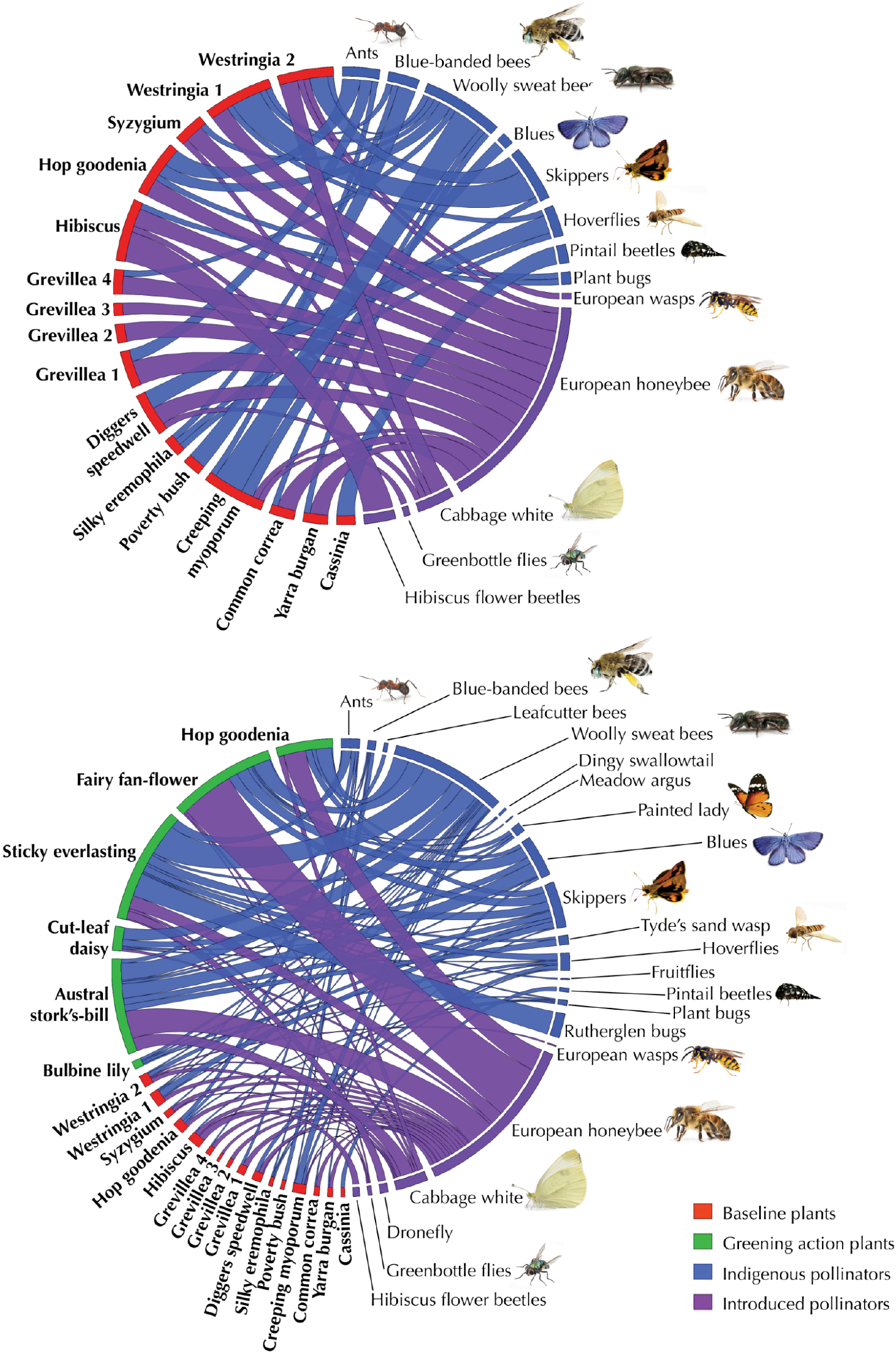
Top: Bipartite quantitative networks of interactions between the baseline plant species (red boxes) and indigenous (blue boxes) and introduced (purple boxes) pollinator and other flower-visiting insect species. Bottom: Bipartite quantitative networks of interactions between the baseline (red Boxes) plus greening action (green boxes) plant species and indigenous (blue boxes) and introduced (purple boxes) pollinator and other flower-visiting insect species. In each network, the width of the chords reflects the frequency of the given interaction as a proportion of the total number of interactions observed.

**Figure 6.**
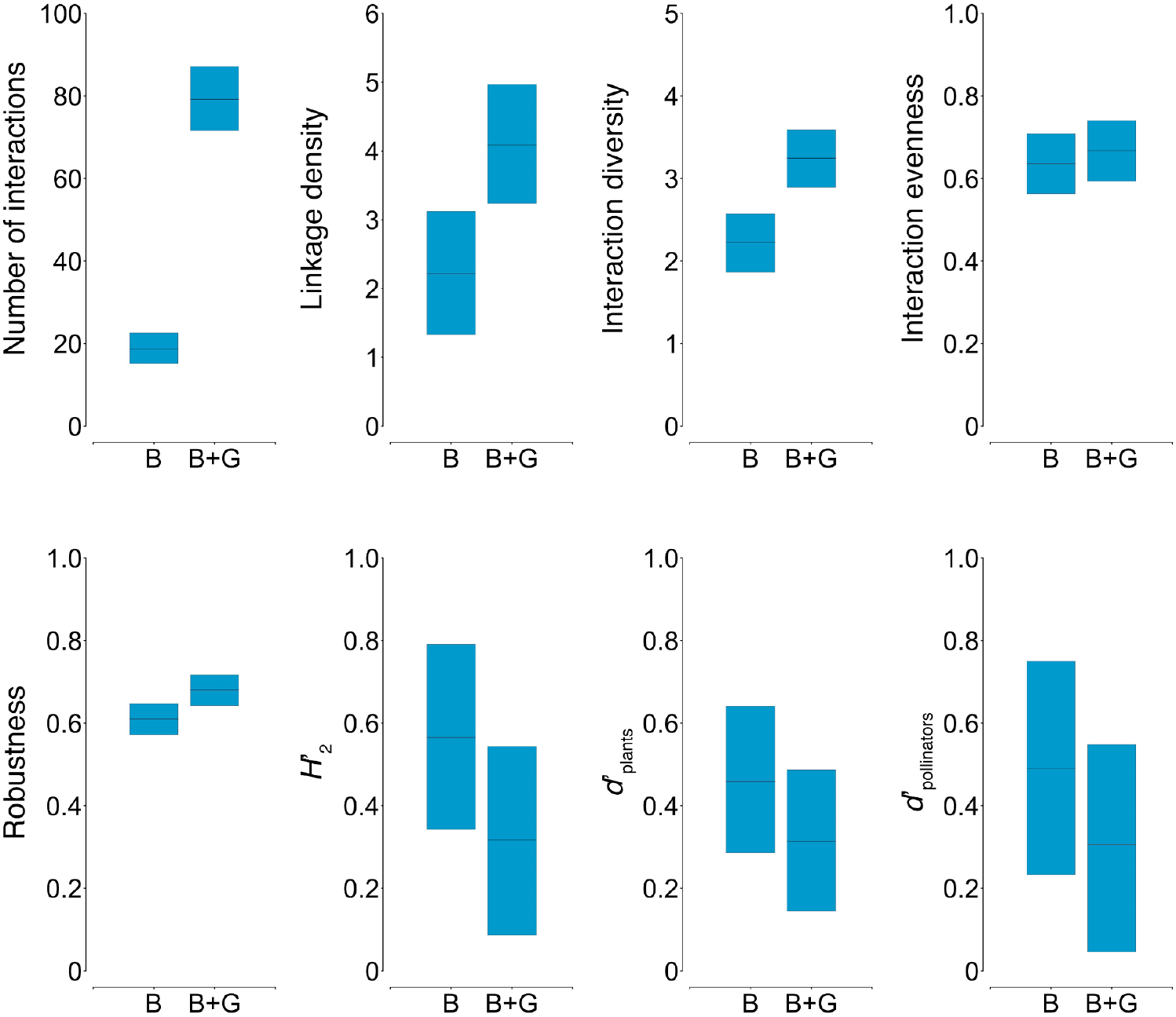
Estimated network-(Number of interactions, Linkage density, Interaction diversity, Interaction evenness, Robustness and network specialisation [*H*_*2*_’]) and species-level (plant and pollinator specialisation [*d*’_plants_ and *d*’_pollinators_]) metrics for the baseline (B) and baseline plus greening action (B+G) plant-pollinator networks. The black horizontal lines represent the mean response and the coloured vertical bars the associated uncertainty (95% credible intervals).

Robustness was one average 1.1 times higher in the greened compared to the baseline network, showing marginally overlapping 95% credible intervals (Table S8; Fig. 6). Network-level specialisation (*H*_*2*_’) and the two species-level metrics of specialisation (*d*’_plants_ and *d*’_pollinators_) were on average lower in the greened compared to the baseline network, showing distinctly overlapping 95% credible intervals (Table S8, Fig. 6).

## Discussion

In this study, we provide empirical evidence of the success of a co-designed practice-research greening action in attracting indigenous pollinators to an urban greenspace. We show how a targeted greening action aimed at partially replacing lawned areas with pollinator-attracting garden beds brought about substantial positive ecological changes in the species occupancy, species richness and network structure of the greenspace’s insect pollinator community. An addition of only six locally indigenous plant species – selected specifically for their known associations with indigenous bees and butterflies – resulted in a 2.5-fold increase in the number of indigenous pollinator species and other flower-visiting insects observed at the study site. The selected indigenous plant species used in the greening action performed distinctly better than the site’s baseline plant species in terms of their capacity to attract indigenous pollinators and contributed to increase the number and diversity of the site’s network of plant-pollinator interactions. As previously advanced by Mata and colleagues (2023), small greening actions, including those carried out in highly urbanised areas, can yield large positive ecological changes across a diverse range of insect functional groups – our findings further contribute to this line of research and practice by providing evidence that small greening actions may also translate into large ecological benefits for insect pollinators and other flower-visiting insects. As such, they support local, regional and global policy aimed at boosting urban sustainability and bolster the argument for increasing the financial support available to fund greening actions for pollinators.

Our findings align closely with a well-established body of empirical research demonstrating the positive effects of urban greenspace floral resource availability on insect pollinator occupancy and species richness (Hennig and Ghazoul 2012, Burdine and McCluney 2019, Wilson and Jamieson 2019, Wenzel et al. 2020, Llodrà-Llabrés and Cariñanos 2022). Similarly, our results lend further support to previous observational (Fukase and Simons 2016, Ballare et al. 2019, Cecala and Wilson Rankin 2021, Mata et al. 2021, Anderson et al. 2023, Zaninotto et al. 2023), experimental (Salisbury et al. 2015, Buchholz and Kowarik 2019, Abbate et al. 2024) and applied research/practice (Brown et al. 2024) works conducted within urban environments highlighting the positive ecological effects of indigenous plants on insect pollinators, particularly locally indigenous ones. While introduced plant species within urban greenspaces also increase the overall quantity and diversity of floral resources available to insect pollinators, they usually establish disproportionally stronger interactions with generalist (often also introduced) insect pollinators, as opposed to with locally, indigenous ones (Zaninotto et al. 2023). Generalist insect pollinators – for example, the European honeybee *Apis mellifera* – are capable of extracting food resources from a diverse array of flower morphologies, thus they are frequently the most dominant and abundant species in plant-pollinator networks within urban environments (Geslin et al. 2013, Zotarelli et al. 2014, Nascimento et al. 2020). Indeed, this species showed the highest frequency of interactions within our study’s plant-pollinator network, both before and after the greening action, and the addition of the greening action indigenous plants increased its overall occupancy across the site.

Importantly however, our data shows that the added indigenous plants established interactions with all the indigenous pollinators observed at the site and that remarkably almost half of these were found interacting exclusively with the new indigenous plants added by the greening action, including a species of leafcutter bee (genus *Megachile*) and three species of butterflies – the dingy swallowtail *Papilio anactus*, the meadow argus *Junonia villida* and the painted lady *Vanessa kershawi*. Additionally, our results indicate that the number of associated indigenous pollinators with a given plant species was on average 3.5 times higher for the added indigenous plants than for the baseline ones and that the probabilities of occurrence of indigenous pollinators were on average invariable higher in the added indigenous than in the baseline plant species. When considered collectively, these findings provide compelling evidence of the success of the investigated co-designed greening action in fulfilling its overarching objective of complementing a targeted urban greenspace with indigenous plant species to support existing and attract new indigenous pollinators and other flower-visiting insects.

The direct modification of the plant community due to the greening action, along with the concomitant changes to the community of insect pollinators, resulted in pronounce changes to the structure of the site’s plant-pollinator network. Notably, our data indicates an average 4.2-fold increase in the number of interactions linking the greened compared to the baseline network, with almost half of these (45%) comprising those between the indigenous plants used in the greening action and indigenous pollinators. This was mirrored by similar but less sharp increases in the density of linkages and diversity of interactions, with no concomitant change to the high evenness of interactions value estimated for the baseline network. In contrast to these diversity metrics, which increase with the number of observations per species and sampling intensity (Kaiser-Bunbury and Blüthgen 2015), the specialisation metrics, which are independent of the completeness of information and can be compared directly across different species within (*d*′) and across (*H*_*2*_′) networks (Kaiser-Bunbury et al. 2017), decreased on average in the greened compared to the baseline network, albeit not to the extent that they may be considered statistically different. This higher functional redundancy and lower specialisation observed in the greened network – indicating increased robustness (Blüthgen and Klein 2011, Kaiser-Bunbury and Blüthgen 2015, as further signalled by the higher values shown by the ‘robustness’ metric – concurs well with previous studies highlighting the positive effects of ecological restoration on disturbed plant–pollinator communities (Kaiser-Bunbury et al. 2017) and complements our earlier findings of the ecological benefits of small greening actions for detritivores, herbivores, predators and parasitoids in urban environments (Mata et al. 2023). In our view, these exciting ecological network results offer compelling evidence of the key role that greening actions – particularly those increasing the floral resources provided by indigenous plants – can play in boosting the capacity of plant-pollinator communities within urban greenspaces to be resilient and maintain ecological functionality in the face of anthropogenic disturbances and climatic fluctuations.

A standout feature of this study was its co-designed methodology, which enabled close alignments between the research questions and methods and the sustainability and biodiversity policy priorities – as well as with the on-ground management realities – of the local government practitioners. Importantly, this approach resonates with calls to unlock the transformative potential of co-design to close the divide between science and practice (Caron et al. 2014, Cadotte et al. 2017, Kurle 2024); as well as with funding directives aimed at assuring that research findings are competently used to inform management practices, strategic planning and policy development. In our study, for example, an initial core priority for the practice team was to work with plant species that were locally indigenous to the municipality – thus closely interwoven with the local Indigenous culture and likely better adapted to thrive in the local soil and climatic conditions (Kawecki and Ebert 2004, Mata et al. 2020, Cumpston et al. 2022) – and known to be closely associated with indigenous bees (primarily blue-banded bees) and butterflies. Notwithstanding this focus, the research team invited the practitioners to consider including in the study other insect taxa – e.g. hoverflies and other flies, ants, heteropteran bugs, beetles and wasps – known for their pollination and other ecological roles in and beyond urban environments (Winfree et al. 2011, Orford et al. 2015, Rader et al. 2015, Silva et al. 2023). This coordinated effort to broaden the study’s taxonomical and functional scope opened the door to a deeper and more nuanced understanding of the rich diversity of pollinators and other flower-visiting insects that may be attracted to urban greenspaces through targeted additions of indigenous plants. Indeed, over half of the species recorded in our study were neither bees nor butterflies, including a fruit fly and a thread-waisted wasp that were recorded exclusively on the greening action plants, both belonging to insect families with many known pollinating species.

We are aware that our study allows for several areas of refinement, paving the way for future research opportunities. To begin with, our richness, occupancy and network metrics estimates were inferred from robustly replicated plant-level data, yet the replication at the site-level was limited to the one greenspace investigated here. Thus, we should sound a note of caution with regards to generalising our findings to other urban systems experiencing a different set of landscape contexts and climatic conditions. Most importantly, however, our data and model estimates can be integrated through metanalytical synthesis with those arising from analogous studies to obtain more comprehensive and generalisable evidence (i.e. where the effects of exogenous drivers are understood) of the relationships between the researched practices and the observed effects (Ockendon et al. 2021). Another shortcoming was the study’s limited focus on food, as opposed to food plus habitat, resources. Beyond the floral resources provided by plants, many insect pollinators – particularly ground-nesting bees such as sweat bees (family Halictidae), which were represented in as much as 18% of the study’s recorded interactions – require habitat for nesting and overwintering (Harrison and Winfree 2015, Threlfall et al. 2015, Schueller et al. 2023). The prospect of being able to overcome the challenge of attracting pollinators and other flower-visiting insects to urban greenspaces through actions that synergistically provides them with both food and habitat resources serves a continuous incentive for future collaborations between researchers and practitioners.

By shedding light on the fact that it possible to replace lawned areas within greenspaces with indigenous midstorey plant species to spark positive ecological outcomes for indigenous pollinators and other flower-visiting insects – including increasing the overall greenspace biodiversity by attracting new indigenous insect species to the site – our findings make a key contribution to the evidence base underpinning best-practise ‘greening for pollinators’ resources and provide encouraging support to sustainability, environment and biodiversity officers across all scales of government, as well as to a wide-ranging group of build-environment practitioners and professionals, responsible for or endeavouring to support existing or bringing insect pollinators back into our urban environments. More generally, our study establishes a pathway and serves as a catalyst for researchers and local government practitioners seeking to work together towards demonstrating that urban greening can be a sound investment in terms of achieving goals from local biodiversity plans, as well as the more overarching mandates enshrined in regional and global sustainability policies. Beyond our findings’ more proximal role in informing on-ground management, greenspace design and ‘Nature in cities’ policy, we draw attention to the added contribution that the greenspace’s strengthen plant and insect community can potentially make in delivering health, well-being, social and cultural benefits to local residents and visitors (Flies et al. 2017, Maller et al. 2018, Lai et al. 2019), including – and perhaps more fundamentally – to children (Davis et al. 2024). Equally importantly, we hope that the study’s planned and resulting emphasis in indigenous species contributes to bring Indigenous culture to the fore, supporting calls to recognise, emphasise and elevate the key core role that Indigenous knowledge has been playing in protecting and caring for nature and in understanding the interwoven relationship between people and nature within and beyond urban environments (Diaz et al. 2018, Mata et al. 2020, Cumpston et al. 2022).

## Authors Contributions

Luis Mata, Sally Dawe, Maree Keenan, Phoenix Wolfe and Amy Hahs co-designed the project; Luis Mata and Estibaliz Palma collected the data; Luis Mata organised and analysed the data; Luis Mata and Estibaliz Palma conceptualised and developed visualisations and tables; Luis Mata, Estibaliz Palma and Amy Hahs interpreted results; Sally Dawe, Maree Keenan and Phoenix Wolfe provided practice and management insights; Luis Mata led the writing of the manuscript; Estibaliz Palma contributed to the writing of the manuscript; Estibaliz Palma, Phoenix Wolfe and Amy Hahs reviewed the manuscript and all authors approved it for publication.

## Acknowledgements

This research was funded by Greater Dandenong City Council. We gratefully acknowledge the contributions provided by Kirstine Oh and the support provided by the on-ground crew members who managed Amersham Reserve and maintained the added indigenous plants garden beds during the study. We acknowledge the Bunurong people of the Kulin Nation as the Traditional Custodians of the lands and waters on which this research took place – we pay our respects to their Elders, past, present and emerging.

## Data availability statement

Data and codes to reproduce models and plots are already published and publicly available in Zenodo: https://doi.org/10.5281/zenodo.14503804

