## Supplementary Information for "Attracting indigenous pollinators to urban greenspaces"

Supporting Information

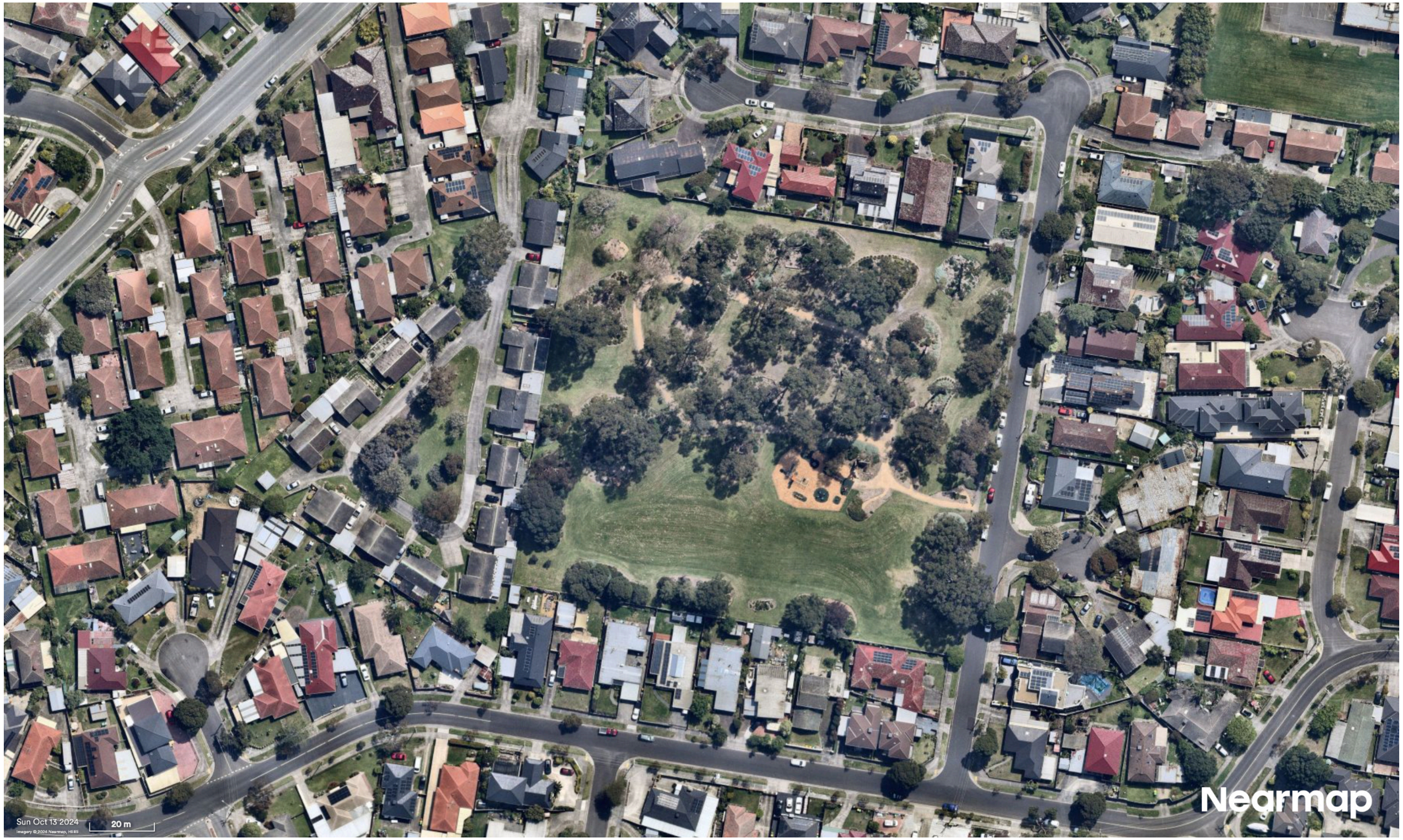


**Figure S1**. Aerial satellite image of Amersham Reserve and its surrounding suburban residential. The greenspace is in the City of Greater Dandenong, Melbourne, Victoria, Australia. Photo provided by Nearmap as accessed through The University of Melbourne's library geospatial resources.

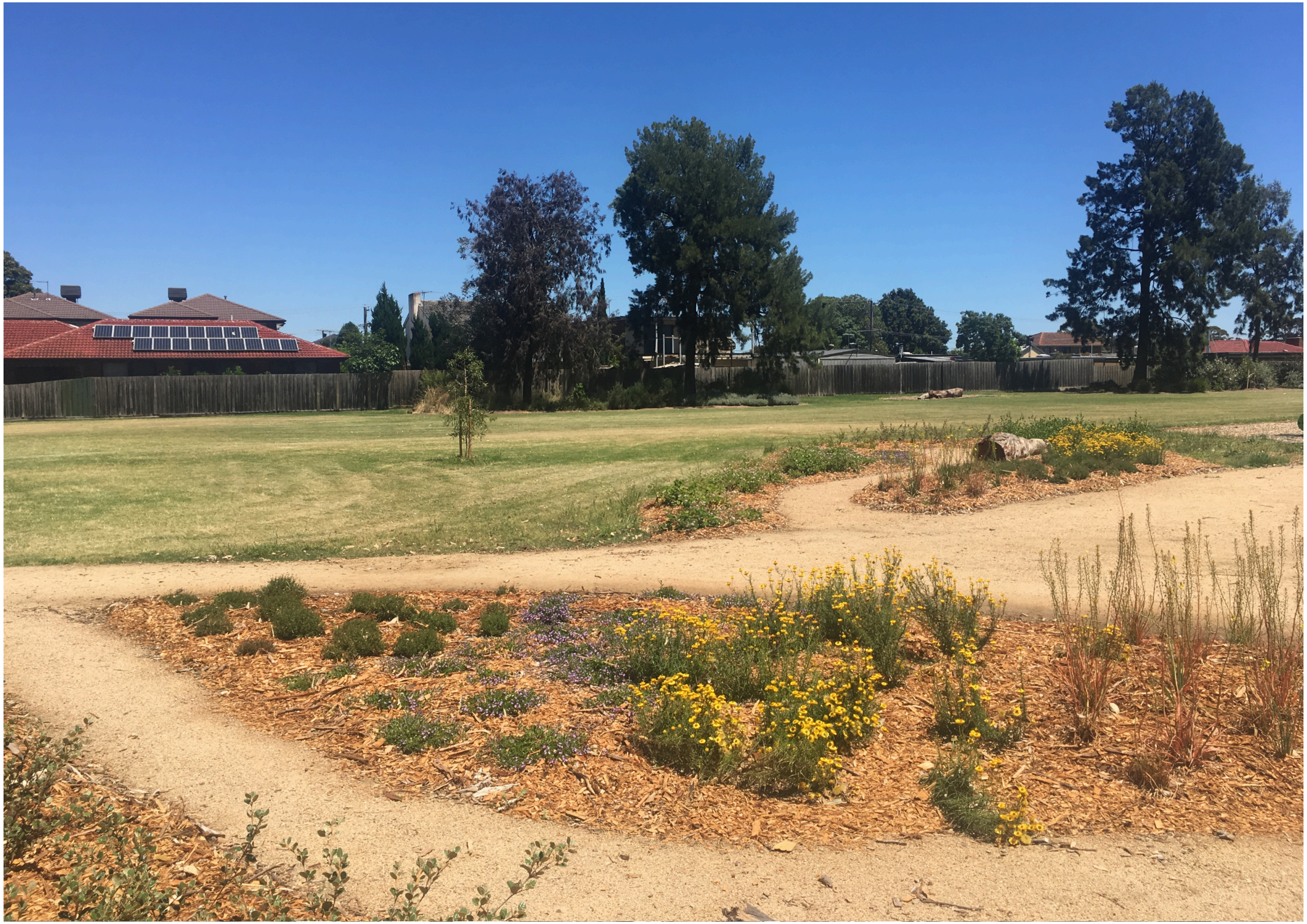

**Figure S2**. Image of the garden beds that were planted in August 2020 with a mix of eight indigenous midstorey plant species, which are referred in the main text as greening action plant species (Table S1). Photo by Luis Mata.

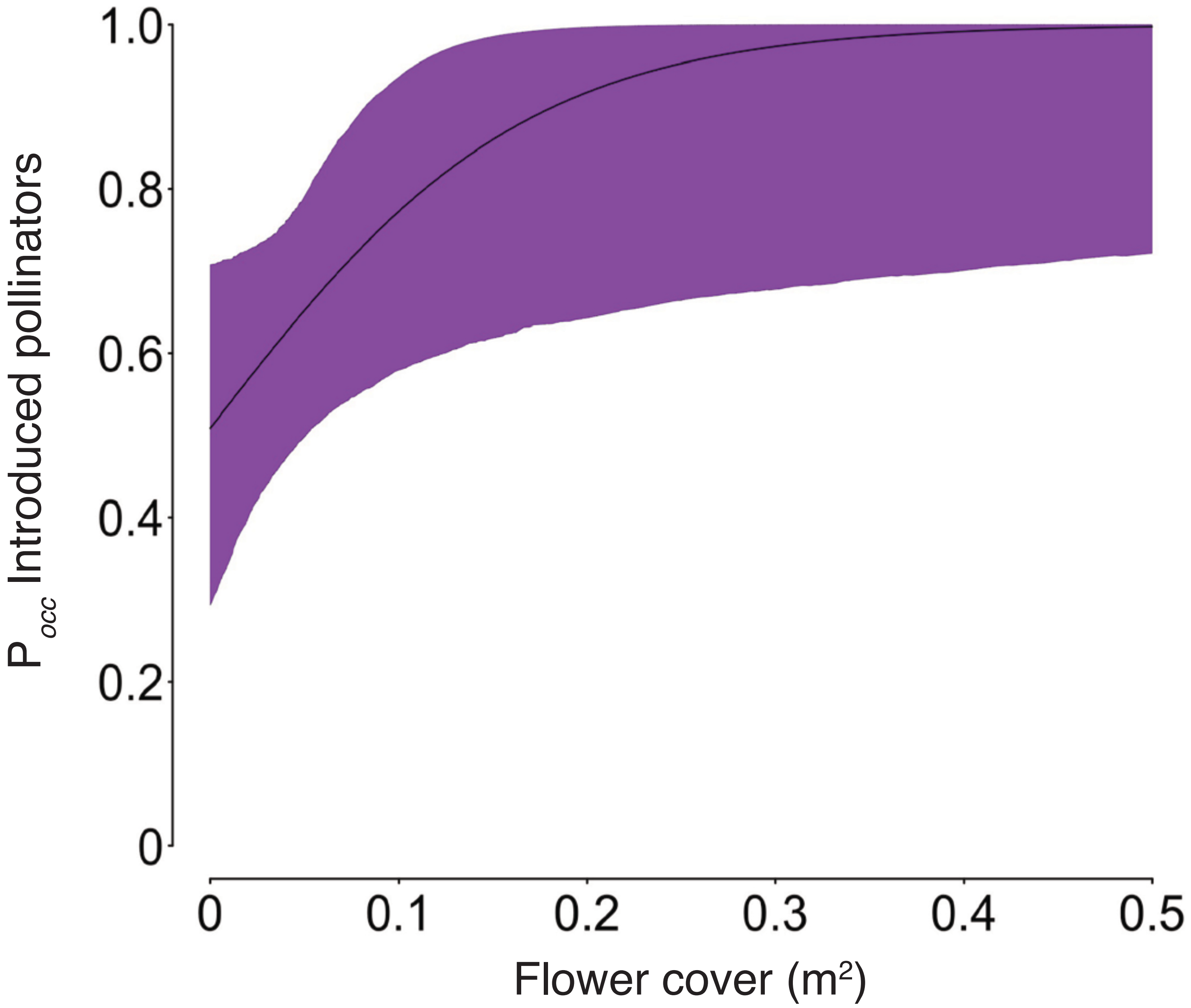


**Figure S3**. Estimated effect of flower cover on the probability of occurrence for introduced pollinator and other flower-visiting insect species. The black curved line represents the mean response and the coloured curved polygon the associated uncertainty (95% credible intervals).

**Table S1**. Taxonomical, biological and ecological information on the 22 plant species that were part of the study.

| Common name | Species | Family | Order | Base clade | Growth form | Origin | Flower colour |
|---|---|---|---|---|---|---|---|
| Baseline plant species | | | | | | | |
| Cassinias | *Cassinia sp. 1* | Asteraceae | Asterales | Superasterids | Shrub | Indigenous | White |
| Common correa | *Correa reflexa* | Rutaceae | Sapindales | Superrosids | Shrub | Indigenous | Red/pink |
| Creeping myoporum | *Myoporum parvifolium* | Scrophulariaceae | Lamiales | Superasterids | Groundcover | Indigenous | White |
| Diggers speedwell | *Veronica perfoliata* | Plantaginaceae | Lamiales | Superasterids | Forb | Native | Blue/violet |
| Grevillea 1 | *Grevillea sp. 1* | Proteaceae | Proteales | Basal eudicots | Shrub | Native | Red/pink |
| Grevillea 2 | *Grevillea sp. 1* | Proteaceae | Proteales | Basal eudicots | Shrub | Native | Red/pink |
| Grevillea 3 | *Grevillea sp. 3* | Proteaceae | Proteales | Basal eudicots | Shrub | Native | Red/pink |
| Grevillea 4 | *Grevillea sp. 4* | Proteaceae | Proteales | Basal eudicots | Shrub | Native | Red/pink |
| Hibiscus 1 | *Hibiscus sp. 1* | Malvaceae | Malvales | Superrosids | Shrub | Introduced | Red/pink |
| Hop goodenia | *Goodenia ovata* | Goodeniaceae | Asterales | Superasterids | Groundcover | Indigenous | Yellow |
| Narrow-leaved poverty bush | *Eremophila alternifolia* | Scrophulariaceae | Lamiales | Superasterids | Shrub | Native | Red/pink |
| Silky eremophila | *Eremophila nivea* | Scrophulariaceae | Lamiales | Superasterids | Shrub | Native | Blue/violet |
| Syzygium 1 | *Syzygium sp. 1* | Myrtaceae | Myrtales | Superrosids | Shrub | Native | White |
| Westringia 1 | *Westringia sp. 1* | Lamiaceae | Lamiales | Superasterids | Shrub | Native | White |
| Westringia 2 | *Westringia sp. 2* | Lamiaceae | Lamiales | Superasterids | Shrub | Native | White |
| Yarra burgan | *Kunzea leptospermoides* | Myrtaceae | Myrtales | Superrosids | Shrub | Indigenous | White |
| Greening action plant species | | | | | | | |
| Austral stork's-bill | *Pelargonium australe* | Geraniaceae | Geraniales | Superrosids | Forb | Indigenous | Red/pink |
| Bulbine lily | *Bulbine bulbosa* | Asphodelaceae | Asparagales | Lilioid monocot | Liliod | Indigenous | Yellow |
| Cut-leaf daisy | *Brachyscome multifida* | Asteraceae | Asterales | Superasterids | Forb | Indigenous | Blue/violet |
| Fairy fan-flower | *Scaevola aemula* | Goodeniaceae | Asterales | Superasterids | Groundcover | Indigenous | Blue/violet |
| Hop goodenia | *Goodenia ovata* | Goodeniaceae | Asterales | Superasterids | Shrub | Indigenous | Yellow |
| Small crowea | *Crowea exalata* | Rutaceae | Sapindales | Superrosids | Shrub | Indigenous | Red/pink |
| Sticky everlasting | *Xerochrysum viscosum* | Asteraceae | Asterales | Superasterids | Forb | Indigenous | Yellow |
| White correa | *Correa alba* | Rutaceae | Sapindales | Superrosids | Shrub | Indigenous | White |

**Table S2**. Origin, flowering time and attractiveness for blue-banded bees (BBB) and butterflies (BUT) of the potential plant species originally considered for the greening action.

| Common name | Scientific name | Family | Origin | Growth form | Flower colour | Sep | Oct | Nov | Dec | Jan | Feb | Mar | Apr | May | Jun | Jul | Aug | BBB | BUT |
|---|---|---|---|---|---|---|---|---|---|---|---|---|---|---|---|---|---|---|---|
| Austral stork's-bill | *Pelargonium australe* | Geraniaceae | Indigenous | Forb | Red/pink | | ■ | ■ | ■ | ■ | ■ | ■ | ■ | ■ | | | | x | x |
| Black-anther flax-lily | *Dianella revoluta* | Asphodelaceae | Indigenous | Lilioid | Blue/violet | ■ | ■ | ■ | ■ | | | | | | | | | x | |
| Blue dampiera | *Dampiera stricta* | Goodeniaceae | Indigenous | Shrub | Blue/violet | ■ | ■ | ■ | ■ | ■ | | | | | | | ■ | | x |
| Bulbine lily | *Bulbine bulbosa* | Asphodelaceae | Indigenous | Lilioid | Yellow | ■ | ■ | ■ | ■ | ■ | ■ | | | | | | | x | |
| Cut-leaf daisy | *Brachyscome multifida* | Asteraceae | Indigenous | Forb | Blue/violet | ■ | ■ | ■ | ■ | ■ | ■ | | | | | | | | x |
| Fairy fan-flower | *Scaveola aemula* | Goodeniaceae | Indigenous | Groundcover | Blue/violet | ■ | ■ | ■ | ■ | ■ | ■ | ■ | ■ | | | | | | x |
| Hop goodenia | *Goodenia ovata* | Goodeniaceae | Indigenous | Shrub | Yellow | ■ | ■ | ■ | ■ | ■ | ■ | ■ | ■ | ■ | ■ | ■ | ■ | x | x |
| River bottlebrush | *Callistemon sieberi* | Myrtaceae | Indigenous | Shrub | White | | | ■ | ■ | ■ | | | | | | | | | x |
| Rounded noon-flower | *Disphyma crassifolium* | Aizoaceae | Indigenous | Groundcover | Red/pink | | ■ | ■ | ■ | ■ | ■ | | | | | | | | x |
| Showy isotome | *Isotoma axilaris* | Campanulaceae | Indigenous | Forb | Blue/violet | ■ | ■ | ■ | ■ | ■ | ■ | ■ | ■ | ■ | ■ | ■ | ■ | x | |
| Small crowea | *Crowea exalata* | Rutaceae | Indigenous | Shrub | Red/pink | ■ | ■ | ■ | ■ | ■ | ■ | | | | | | | x | |
| Sticky everlasting | *Xerochrysum viscosum* | Asteraceae | Indigenous | Forb | Yellow | ■ | ■ | ■ | ■ | ■ | ■ | ■ | | | | | | | x |
| Twiggy daisy-bush | *Olearia ramulosa* | Asteraceae | Indigenous | Shrub | White | | ■ | ■ | ■ | ■ | ■ | ■ | ■ | ■ | | | | | x |
| Yarra burgan | *Kunzea leptospermoides* | Myrtaceae | Indigenous | Shrub | White | ■ | ■ | ■ | ■ | | | | | | | | | | x |

**Table S3**. Taxonomical and ecological information on the insect pollinator and other flower-visiting insect species observed at Amersham Reserve during the study.

| Order | Superfamily | Species/morphospecies | Common name | Family | Origin |
|---|---|---|---|---|---|
| Coleoptera | *Cucujoidea* | Aethina concolor | Hibiscus flower beetles | Nitidulidae | Introduced |
| | *Tenebrionoidea* | Mordellidae 1 | Pintail beetles | Mordellidae | Indigenous |
| Diptera | *Oestroidea* | Lucilia sp. 1 | Greenbottle flies | Calliphoridae | Introduced |
| | *Tephritoidea* | Tephritidae 1 | Fruit flies | Tephritidae | Indigenous |
| | *Syrphoidea* | Eristalis tenax | Dronefly | Syrphidae | Introduced |
| | | Syrphidae 1 | Hoverflies | Syrphidae | Indigenous |
| Hemiptera | *Lygaeoidea* | Nysius sp. 1 | Rutherglen bugs | Lygaeidae | Indigenous |
| | *Miroidea* | Miridae 1 | Plant bugs | Miridae | Indigenous |
| Hymenoptera | *Apoidea* | Amegilla sp. 1 | Blue-banded bees | Apidae | Indigenous |
| | | Apis mellifera | European honeybee | Apidae | Introduced |
| | | Lasioglossum sp. 1 | Wolly sweat bees | Halictidae | Indigenous |
| | | Megachile sp. 1 | Leafcutter bees | Megachilidae | Indigenous |
| | | Podalonia tydei | Tyde's sand wasp | Sphecidae | Indigenous |
| | *Formicoidea* | Formicidae 1 | Ants | Formicidae | Indigenous |
| | *Vespoidea* | Vespula sp. 1 | European wasps | Vespidae | Introduced |
| Lepidoptera | *Papilionoidea* | Hesperiidae 1 | Skippers | Hesperiidae | Indigenous |
| | | Junonia villida | Meadow argus | Nymphalidae | Indigenous |
| | | Lycaneidae 1 | Blues | Lycaneidae | Indigenous |
| | | Papilio anactus | Dingy swallowtail | Papilionidae | Indigenous |
| | | Pieris rapae | Cabbage white | Pieridae | Introduced |
| | | Vanessa kershawi | Painted lady | Nymphalidae | Indigenous |

**Table S4**. Number of records of each pollinator and other flower-visiting insect species observed on each plant species during the study.

| | Ants | Blue-banded bees | Blues | Cabbage white | Dingy swallowtail | Dronefly | European honeybee | European wasps | Fruit flies | Greenbottle flies | Hibiscus flower beetle | Hoverflies | Leafcutter bees | Meadow argus | Painted lady | Pintail beetles | Plant bugs | Rutherglen bugs | Skippers | Tyde's sand wasp | Woolly sweat bees |
|---|---|---|---|---|---|---|---|---|---|---|---|---|---|---|---|---|---|---|---|---|---|
| Austral stork's-bill | 0 | 0 | 11 | 10 | 1 | 0 | 21 | 0 | 0 | 0 | 0 | 0 | 3 | 0 | 2 | 0 | 0 | 0 | 5 | 5 | 13 |
| Bulbine lily | 0 | 0 | 1 | 0 | 0 | 0 | 0 | 0 | 0 | 0 | 0 | 2 | 0 | 0 | 0 | 0 | 1 | 0 | 0 | 0 | 2 |
| Cassinias | 0 | 0 | 0 | 0 | 0 | 0 | 0 | 0 | 0 | 0 | 0 | 0 | 0 | 0 | 0 | 3 | 0 | 0 | 0 | 0 | 0 |
| Common correa | 0 | 0 | 0 | 0 | 0 | 0 | 3 | 0 | 0 | 0 | 0 | 0 | 0 | 0 | 0 | 0 | 0 | 0 | 0 | 0 | 1 |
| Creeping myoporum | 1 | 0 | 2 | 0 | 0 | 0 | 1 | 0 | 0 | 1 | 0 | 0 | 0 | 0 | 0 | 0 | 0 | 0 | 0 | 0 | 5 |
| Cut-leaf daisy | 0 | 0 | 5 | 0 | 0 | 0 | 2 | 0 | 0 | 0 | 0 | 1 | 0 | 0 | 0 | 0 | 0 | 0 | 1 | 0 | 9 |
| Diggers speedwell | 0 | 0 | 0 | 1 | 0 | 0 | 3 | 0 | 0 | 0 | 0 | 3 | 0 | 0 | 0 | 0 | 0 | 0 | 0 | 0 | 0 |
| Fairy fan-flower | 1 | 0 | 9 | 2 | 1 | 0 | 41 | 0 | 0 | 1 | 0 | 0 | 0 | 1 | 3 | 0 | 0 | 0 | 7 | 0 | 13 |
| Grevillea 1 | 2 | 0 | 0 | 0 | 0 | 0 | 4 | 0 | 0 | 0 | 0 | 0 | 0 | 0 | 0 | 0 | 0 | 0 | 0 | 0 | 0 |
| Grevillea 2 | 0 | 0 | 0 | 0 | 0 | 0 | 3 | 0 | 0 | 0 | 0 | 0 | 0 | 0 | 0 | 0 | 0 | 0 | 0 | 0 | 0 |
| Grevillea 3 | 0 | 0 | 0 | 0 | 0 | 0 | 2 | 0 | 0 | 0 | 0 | 0 | 0 | 0 | 0 | 0 | 0 | 0 | 0 | 0 | 0 |
| Grevillea 4 | 0 | 0 | 0 | 0 | 0 | 0 | 3 | 0 | 0 | 0 | 0 | 0 | 0 | 0 | 0 | 0 | 0 | 0 | 0 | 0 | 1 |
| Hibiscus 1 | 0 | 0 | 0 | 1 | 0 | 0 | 3 | 0 | 0 | 0 | 5 | 0 | 0 | 0 | 0 | 0 | 1 | 0 | 0 | 0 | 0 |
| Hop goodenia | 0 | 1 | 3 | 4 | 0 | 0 | 17 | 0 | 0 | 0 | 0 | 0 | 0 | 0 | 0 | 0 | 0 | 0 | 5 | 0 | 12 |
| Hop goodenia gc | 3 | 0 | 0 | 0 | 0 | 0 | 3 | 0 | 0 | 0 | 0 | 0 | 0 | 0 | 0 | 0 | 1 | 0 | 0 | 0 | 2 |
| Narrow-leaved poverty bush | 0 | 3 | 0 | 0 | 0 | 0 | 0 | 0 | 0 | 0 | 0 | 0 | 0 | 0 | 0 | 0 | 0 | 0 | 0 | 0 | 0 |
| Silky eremophila | 0 | 1 | 0 | 0 | 0 | 0 | 0 | 0 | 0 | 0 | 0 | 1 | 0 | 0 | 0 | 0 | 0 | 0 | 1 | 0 | 0 |
| Sticky everlasting | 7 | 0 | 0 | 2 | 0 | 6 | 10 | 0 | 1 | 1 | 0 | 5 | 0 | 0 | 2 | 0 | 1 | 18 | 7 | 2 | 24 |
| Syzygium 1 | 0 | 0 | 0 | 1 | 0 | 0 | 3 | 0 | 0 | 0 | 0 | 0 | 0 | 0 | 0 | 0 | 0 | 0 | 0 | 0 | 1 |
| Westringia 1 | 0 | 1 | 0 | 0 | 0 | 0 | 3 | 0 | 0 | 0 | 0 | 1 | 0 | 0 | 0 | 0 | 0 | 0 | 5 | 0 | 1 |
| Westringia 2 | 0 | 0 | 0 | 3 | 0 | 0 | 1 | 1 | 0 | 0 | 0 | 0 | 0 | 0 | 0 | 0 | 0 | 0 | 3 | 0 | 1 |
| Yarra burgan | 0 | 0 | 0 | 0 | 0 | 0 | 3 | 0 | 0 | 0 | 0 | 0 | 0 | 0 | 0 | 0 | 0 | 0 | 0 | 0 | 1 |

**Table S5**. Posterior estimates of the species richness of pollinator and other flower-visiting insect species for the baseline and greening action plant species, and for each individual plant species. SD: Standard deviation; CI: Credible interval.

| Species richness of indigenous pollinators | Mean | SD | CI: 2.5% | CI:97.5% |
|---|---|---|---|---|
| By plant group | | | | |
| Baseline plant species | 1.769 | 0.148 | 1.500 | 2.125 |
| Greening action plant species | 6.140 | 0.303 | 5.833 | 6.833 |
| By plant species | | | | |
| Austral stork's-bill | 7.236 | 0.462 | 7 | 8 |
| Bulbine lily | 4.009 | 0.097 | 4 | 4 |
| Cassinias | 1.000 | 0.000 | 1 | 1 |
| Common correa | 1.285 | 0.460 | 1 | 2 |
| Creeping myoporum | 3.715 | 0.726 | 3 | 5 |
| Cut-leaf daisy | 4.778 | 0.999 | 4 | 7 |
| Diggers speedwell | 1.397 | 0.493 | 1 | 2 |
| Fairy fan-flower | 7.249 | 0.473 | 7 | 8 |
| Grevillea 1 | 1.052 | 0.222 | 1 | 2 |
| Grevillea 2 | 0.218 | 0.413 | 0 | 1 |
| Grevillea 3 | 0.000 | 0.000 | 0 | 0 |
| Grevillea 4 | 1.035 | 0.184 | 1 | 2 |
| Hibiscus 1 | 1.363 | 0.563 | 1 | 3 |
| Hop goodenia | 4.191 | 0.414 | 4 | 5 |
| Hop goodenia gc | 3.330 | 0.739 | 3 | 5 |
| Narrow-leaved poverty bush | 1.027 | 0.163 | 1 | 2 |
| Silky eremophila | 3.792 | 0.735 | 3 | 5 |
| Sticky everlasting | 9.379 | 0.995 | 9 | 12 |
| Syzygium 1 | 1.274 | 0.469 | 1 | 2 |
| Westringia 1 | 4.356 | 0.544 | 4 | 6 |
| Westringia 2 | 2.453 | 0.555 | 2 | 4 |
| Yarra burgan | 1.002 | 0.049 | 1 | 1 |

**Table S6**. Posterior estimates of the probabilities of occurrence of indigenous and introduced insect pollinator species for the baseline and greening action plant species. SD: Standard deviation; CI: Credible interval.

| Probability of occurrence of insect pollinators | Baseline plant species | | | | Greening action plant species | | | |
|---|---|---|---|---|---|---|---|---|
| | **Mean** | **SD** | **CI: 2.5%** | **CI: 97.5%** | **Mean** | **SD** | **CI: 2.5%** | **CI: 97.5%** |
| Indigenous species | | | | | | | | |
| Blue-banded bees | 0.183 | 0.142 | 0.035 | 0.579 | 0.388 | 0.197 | 0.112 | 0.846 |
| Blues | 0.205 | 0.057 | 0.109 | 0.331 | 0.458 | 0.082 | 0.303 | 0.626 |
| Hoverflies | 0.123 | 0.059 | 0.045 | 0.270 | 0.310 | 0.107 | 0.145 | 0.564 |
| Skippers | 0.170 | 0.049 | 0.088 | 0.279 | 0.402 | 0.077 | 0.260 | 0.560 |
| Woolly sweat bees | 0.404 | 0.078 | 0.264 | 0.565 | 0.690 | 0.069 | 0.548 | 0.819 |
| Introduced species | | | | | | | | |
| Cabbage white | 0.213 | 0.076 | 0.101 | 0.395 | 0.466 | 0.107 | 0.277 | 0.700 |
| European honeybee | 0.403 | 0.070 | 0.273 | 0.546 | 0.690 | 0.065 | 0.558 | 0.809 |

**Table S7**. Posterior estimates of the effects of flower cover on the probabilities of occurrence of indigenous and introduced pollinator and other flower-visiting insect species. SD: Standard deviation; CI: Credible interval.

| **Model parameter** | **Mean** | **SD** | **CI: 2.5%** | **CI: 97.5%** |
|---|---|---|---|---|
| Indigenous species | | | | |
| Pr (occurrence) | 0.933 | 0.049 | 0.816 | 0.994 |
| Pr (detection) | 0.617 | 0.040 | 0.537 | 0.694 |
| Model intercept, a | 2.955 | 0.938 | 1.490 | 5.063 |
| Effect size, b | 3.567 | 1.733 | 0.859 | 7.416 |
| Introduced species | | | | |
| Pr (occurrence) | 0.795 | 0.106 | 0.603 | 0.973 |
| Pr (detection) | 0.724 | 0.046 | 0.631 | 0.809 |
| Model intercept, a | 1.546 | 0.857 | 0.418 | 3.591 |
| Effect size, b | 2.220 | 1.633 | 0.190 | 6.048 |

**Table S8**. Posterior estimates for the network- and species-level metrics calculated for the baseline and baseline plus greening action networks. SD: Standard deviation; CI: Credible interval.

| **Network metrics** | **Baseline network** | | | | **Baseline + greening action network** | | | |
|---|---|---|---|---|---|---|---|---|
| | **Mean** | **SD** | **CI: 2.5%** | **CI: 97.5%** | **Mean** | **SD** | **CI: 2.5%** | **CI: 97.5%** |
| Number of interactions | 18.640 | 1.883 | 15.192 | 22.429 | 79.054 | 4.088 | 71.121 | 87.100 |
| Linkage density | 2.215 | 0.456 | 1.314 | 3.146 | 4.099 | 0.464 | 3.164 | 5.021 |
| Interaction diversity | 2.221 | 0.173 | 1.866 | 2.561 | 3.247 | 0.171 | 2.911 | 3.601 |
| Interaction evenness | 0.636 | 0.037 | 0.562 | 0.708 | 0.668 | 0.037 | 0.596 | 0.739 |
| Robustness | 0.614 | 0.020 | 0.573 | 0.656 | 0.675 | 0.020 | 0.635 | 0.715 |
| $H'_2$ | 0.566 | 0.113 | 0.337 | 0.790 | 0.314 | 0.111 | 0.095 | 0.536 |
| $d'_{plants}$ | 0.458 | 0.083 | 0.290 | 0.625 | 0.313 | 0.085 | 0.147 | 0.487 |
| $d'_{pollinators}$ | 0.491 | 0.134 | 0.230 | 0.755 | 0.304 | 0.133 | 0.048 | 0.557 |
